# Inferring time-derivatives, including cell growth rates, using Gaussian processes

**DOI:** 10.1101/055483

**Authors:** Peter S. Swain, Keiran Stevenson, Allen Leary, Luis F. Montano-Gutierrez, Ivan B. N. Clark, Jackie Vogel, Teuta Pilizota

## Abstract

Often the time-derivative of a measured variable is of as much interest as the variable itself. For a growing population of biological cells, for example, the population's growth rate is typically more important than its size. Here we introduce a non-parametric method to infer first and second time-derivatives as a function of time from time-series data. Our approach is based on established properties of Gaussian processes and therefore applies to a wide range of data. In tests, the method is at least as accurate as others, but has several advantages: it estimates errors both in the inference and in any summary statistics, such as lag times, allows interpolation with the corresponding error estimation, and can be applied to any number of experimental replicates. As illustrations, we infer growth rate from measurements of the optical density of populations of microbial cells and estimate the rate of *in vitro* assembly of an amyloid fibril and both the speed and acceleration of two separating spindle pole bodies in a single yeast cell. Being accessible through both a GUI and from scripts, our algorithm should have broad application across the sciences.

---

Estimating the time-derivatives of a signal is a common task in science. A well-known example is the growth rate of a population of cells, which is defined as the time-derivative of the logarithm of the population size [1] and is used extensively in both the life sciences and biotechnology.

A common approach to estimate such derivatives is to fit a mathematical equation that, say, describes cellular growth and so determine the maximum growth rate from the best-fit value of a parameter in the equation [2]. Such parametric approaches rely, however, on the mathematical model being a suitable description of the underlying biological or physical process, and, at least for cellular growth, it is common to find examples where the standard models are not appropriate [3].

The alternative is to use a non-parameteric method and so estimate time-derivatives directly from the data. Examples include taking numerical derivatives [4] or using local polynomial or spline estimators [5]. Although these approaches do not require knowledge of the underlying process, it can be difficult to determine the error in their estimation [5] and to incorporate experimental replicates, which with wide access to high throughput technologies, are now the norm.

Here we develop a methodology that uses Gaussian processes to fit time-series and infer both the first and second time-derivatives. One advantage of using Gaussian processes over parametric approaches is that we can fit a wider variety of data. Rather than assuming that a particular function characterises the data (a particular mathematical equation), we instead make assumptions about the family of functions that can describe the data. An infinite number of functions exist in this family, and the family can capture many more temporal trends in the data than any one equation. The advantages over existing non-parametric methods are that we can straightforwardly and systematically combine data from replicate experiments (by simply pooling all datasets) and predict errors both in the estimations of derivatives and in any summary statistics. A potential disadvantage because we use Gaussian processes is that we must assume that the measurement noise has a normal or log-normal distribution (as do many other methods), but we can relax this assumption if there are multiple experimental replicates.

To illustrate how our approach predicts errors and can combine information from experimental replicates, we first focus on inferring growth rate from measurements of the optical density of a growing population of biological cells. Plate readers, which are now wide-spread, make such data easy to obtain, typically with hundreds of measurements and often at least 3–10 replicates. We will also, though, show other examples: estimating the rate of *in vitro* assembly of an amyloid fibril and inferring the speed and acceleration of two separating spindle pole bodies in a single yeast cell.

## Results

To verify our algorithm,s inference of first and second time-derivatives, we followed the tests of De Brabanter *et al.* [6]. Gaussian measurement noise was added to an analytic function for which time-derivatives can be found exactly, and the mean absolute difference between the inferred derivative and the exact derivative was used to score the inference (see [6] for details - the end points are not included). We show the distribution of scores for 100 different datasets each with a different sample of the measurement noise (Fig. 1).

**Figure 1.**
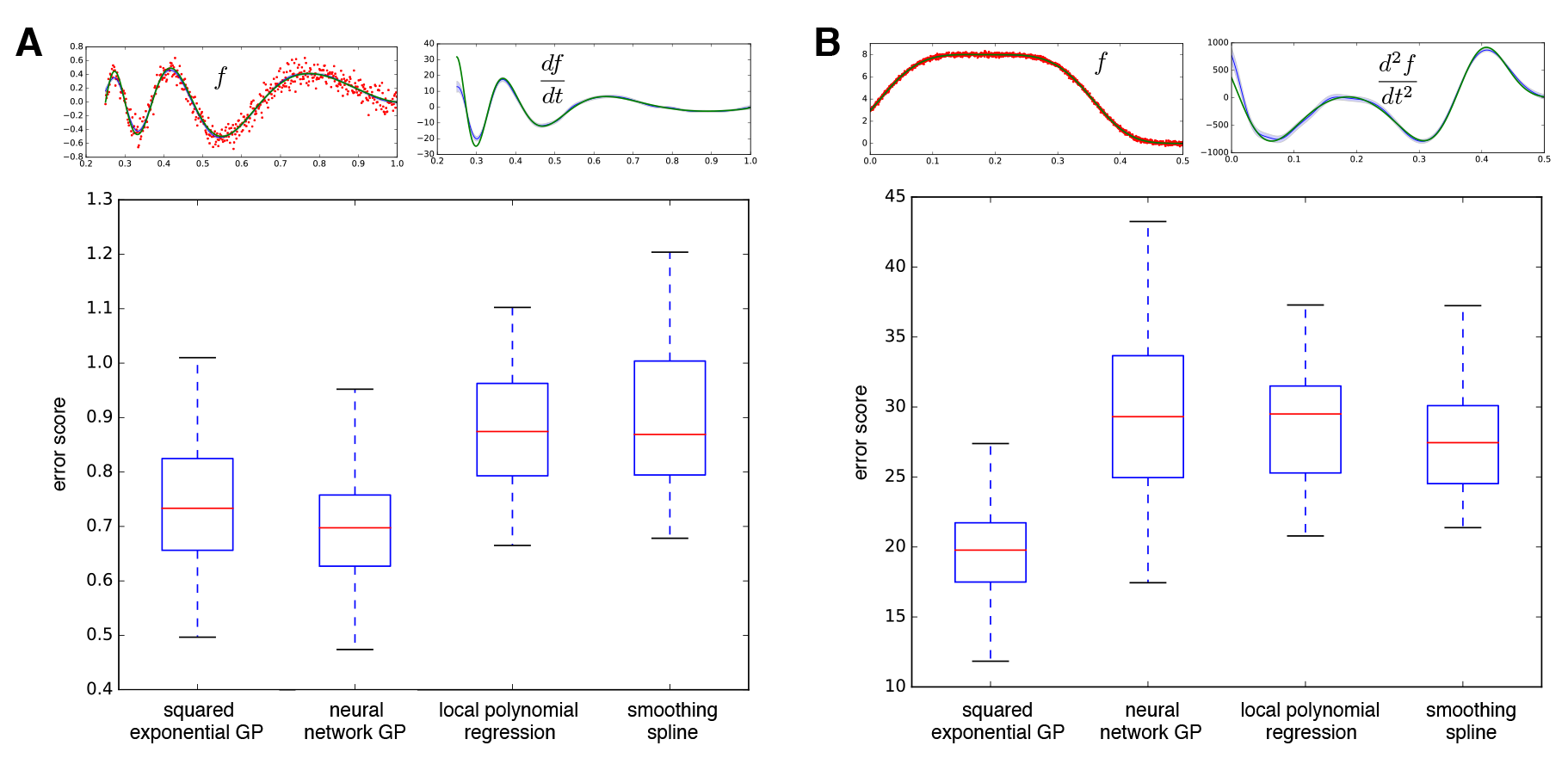
The inference method can perform better than alternatives. A) Inference of the first derivative. A box plot of error scores (related to the mean absolute difference between the inferred and exact derivative) for inference of the first derivative. We use either a squared exponential covariance function or a neural network covariance function for our Gaussian process (GP) and compare with local polynomial regression (with *p* = 3) and a quintic penalised smoothing spline (data for both from 6). Top left shows one sample data set (in red with 500 data points), the true underlying function (in green), and the inferred latent function using a neural network covariance function - the best fit (in blue); top right shows the corresponding first derivative (with here an error score of 0.64): exact (in green) and inferred (in blue). Errors (in light blue) are standard deviations. B) Inference of the second derivative. A box plot of scores for inference of the second derivative. The two alternatives are local polynomial regression (with *p* = 5) and a septic penalised smoothing spline (data for both from 6). Top right shows one sample data set (in red with 1500 data points), the underlying function (in green), and the inferred latent function using a neural network covariance function (in blue); top left shows the corresponding second derivative (with here an error score of 26.2): exact (in green) and inferred (in blue).

For these tests, our method outperforms established alternatives. Using Gaussian processes, we specify the family of functions that we consider for our inference by choosing a covariance function (Methods). The result of the fitting is to obtain a probability distribution for the so-called latent function, the function that underlies the data. We report the mean of this distribution as the best-fit latent function. For illustration, we use two covariance functions: a squared exponential covariance function, which only imposes that the functions are smooth over some length scale determined from the data, and a neural network covariance function, which can generate approximately sigmoidal functions[7]. Independent of the choice of covariance function, the method performed at least as well as alternatives (Fig. 1). For most datasets, we find, however, that the squared exponential covariance function is the best choice because it makes the least restrictive assumptions.

We now turn to inferring microbial growth rates (strictly, we infer the specific growth rate: the time derivative of the logarithm of the population size). We fit optical densities, which are proportional to the number of cells if properly calibrated [8], and show that we can infer growth rates for two cases that cannot be easily described by published growth equations [3]. The first exhibits a diaxuic shift with two distinct phases of growth and the second shows an exceptionally long lag (Fig. 2). We infer the growth rate and the estimated errors in our inference as a function of time using all experimental replicates. Data from replicate measurements are pooled together, and the algorithm applied as for a single replicate.

**Figure 2.**
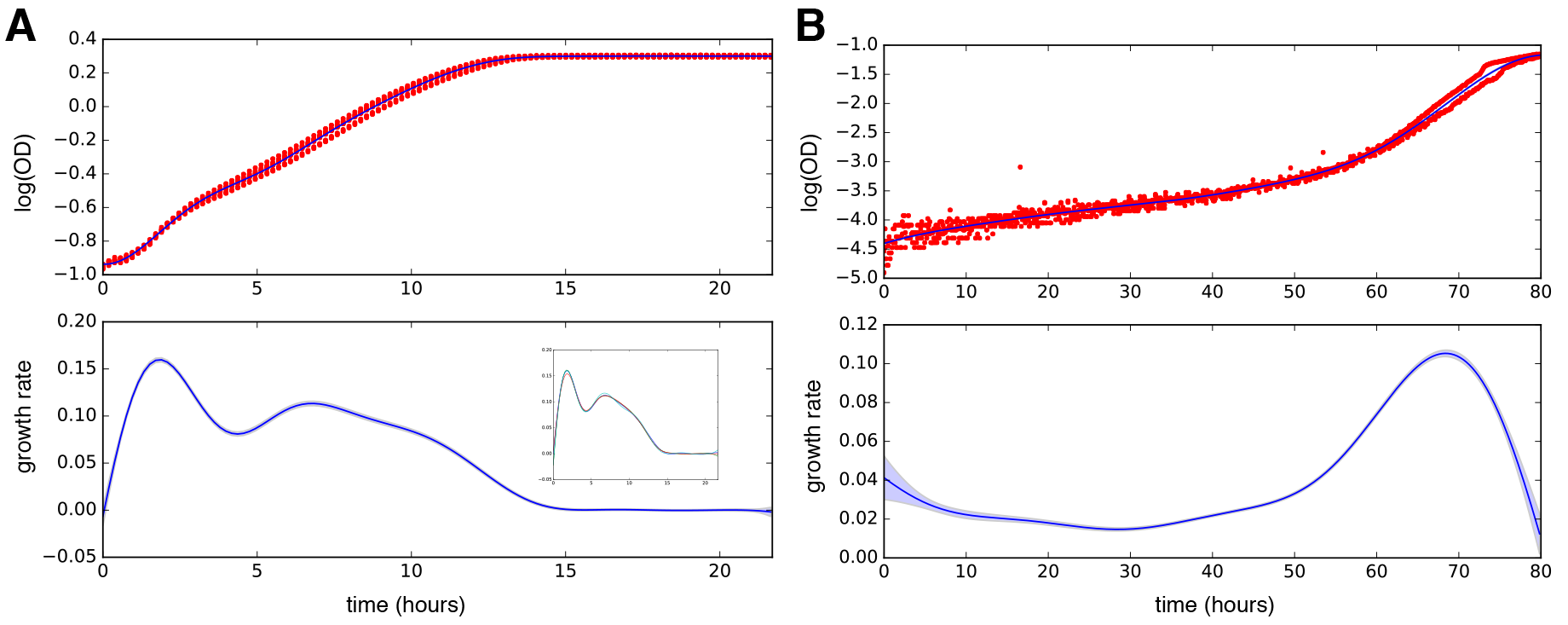
Microbial growth rates can be inferred as a function of time. A) A growth curve of *Saccharomyces cerevisiae* in a mixture of 0.4% glucose and 1% galactose showing a diaxuic shift (7 replicates). The best-fit (mean) latent function is shown in dark blue and the inferred growth rate is shown below. All error bars (light blue) are standard deviations. The inset shows, as an example, 4 sample estimates of the growth rate as a function of time (samples of the first derivative of the latent function - the corresponding samples of the latent function itself are not shown). B) Growth of *Escherichia coli* in hyperosmotic conditions with an unusually long lag and short growth period (2 replicates) and the inferred growth rate. Error bars (light blue) are standard deviations.

Having the inferred growth rate over time can make identifying different stages of the growth curve substantially easier than making this identification from the optical density data alone. For example, the local minimum in the growth rate of Fig. 2A is expected to indicate a shift from cells using glucose to using galactose. Inferring a time-dependent growth rate should increase the robustness of high throughput automated studies, which usually focus on identifying exponential growth [9, 10].

Often summary statistics are used to describe a growth curve, such as the maximum growth rate and the lag time [2], and we can estimate such statistics and their associated errors. From our inference, we find a probability distribution of latent functions that are consistent with the data (Methods). For the best-fit, we report the mean of this distribution, but we can also sample latent functions from the distribution. Each sample provides an example of a latent function that ‘fits, the data. To estimate errors in statistics, we generate say 100 samples of the latent function and its time-derivatives (Fig. 2A inset). For each sample, we calculate the statistic for that sample (for example, the maximum growth rate). We therefore obtain a probability distribution for the statistic and report the mean and standard deviation of this distribution as the best-fit value and the estimated error (0.16 ± 0.002 hr^−1^ for the maximum growth rate for the data in Fig. 2A). A similar approach applies for any statistic that can be calculated from a single growth curve (Methods).

The data for Fig. 2B are considerably noisier than the data for Fig. 2A, and the spread of data is larger at short times than at long times. The magnitude of the measurement error changes with time. More correctly, we typically assume that the measurement error can be described by a Gaussian distribution with zero mean and a constant standard deviation. The magnitude of the measurement error is this standard deviation, and the standard deviation here, for the data of Fig. 2B, appears to change with time (it is largest at early times). To empirically estimate the relative scale of this change, we calculate the variance across replicates at each time point. We assume that the magnitude of the measurement error is a time-independent constant multiplied by this time-dependent relative scale, and we fit that constant (Methods).

As additional examples, we first infer the rate of assembly of an amyloid fibril as a function of time from *in vitro* data (Fig. 3A) [11]. Despite each replicate having high measurement noise compared to the microbial data, the rate of fibril assembly can be inferred accurately because of the many replicates. The second example is one where both the first and the second derivative are useful: estimating the speed of separation of the spindle poles during anaphase (Fig. 3B). We demonstrate that we can infer both time-derivatives and their errors from a single replicate. As expected, the size of the estimated error increases for the first derivative relative to the error in the regression and increases again for the second derivative. Changes in the speed of separation (extrema in the second derivative) are used to characterise anaphase [12] into the fast, pause and slow elongation phases [13]. We chose a Gaussian process with a neural network covariance function for this data rather than the squared exponential covariance function used for the others. The latent functions generated then tend to be flatter either side of the increase in separation: a difference that is important here because we only have a single replicate and that leads to smoother inferences of the acceleration.

**Figure 3.**
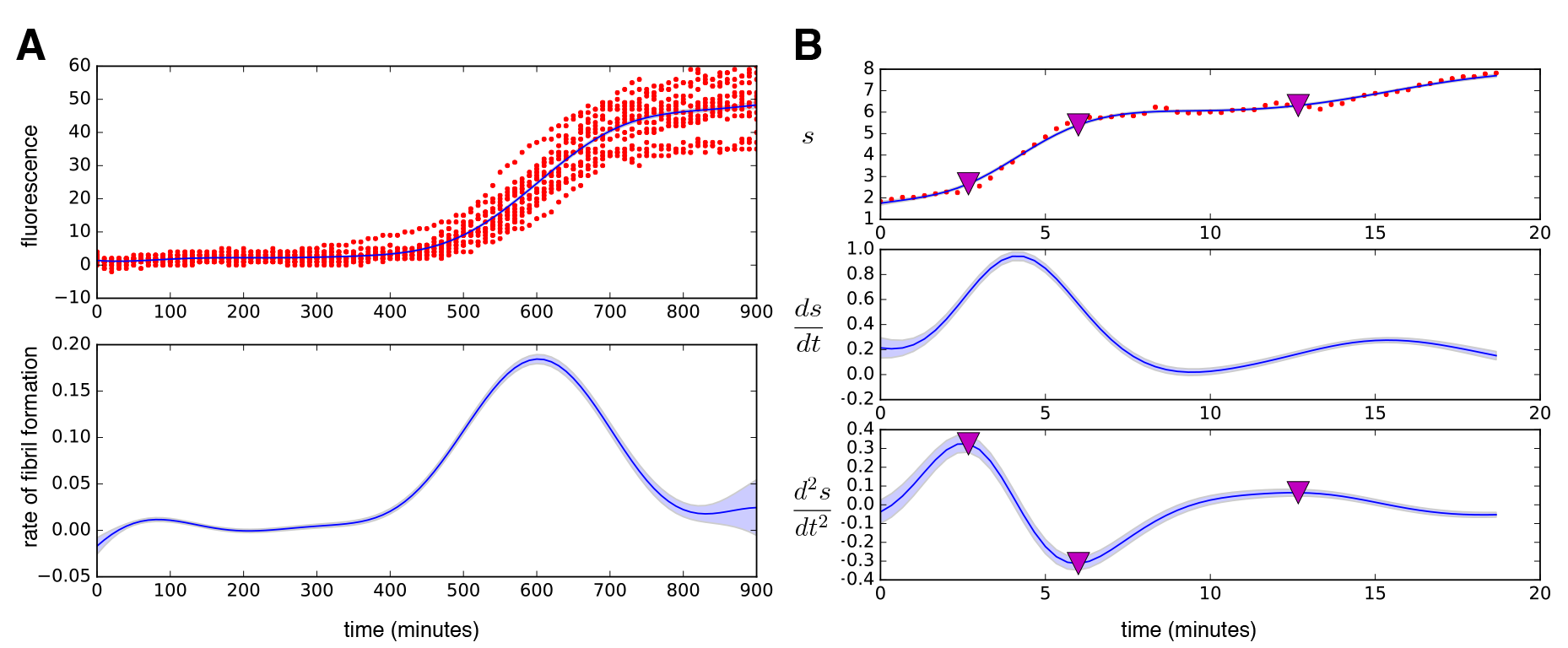
The algorithm has wide application. A) Inferring the *in vitro* rate of assembly of an amyloid fibril. Fluorescence data reporting the formation of fibrils in bovine insulin (at a concentration of 0.1 mg ml^−1^) by the binding of the dye Thioflavin T are shown (red dots) with 15 replicates [11]. The best-fit (top) and the inferred rate of fibril assembly (bottom) are shown in dark blue. Errors (in light blue) are standard deviations. B) Inferring the speed and acceleration of separation of the spindle poles in *S. cerevisiae.* The distance, s, between the two spindles in a single cell is plotted in microns as a function of time (red dots). The best-fit and the inferred speed (middle) and acceleration (bottom) are shown in dark blue. The purple triangles denote turning points in the acceleration and separate anaphase into stages with fast and slow elongation separated by a pause [12]. Errors are standard deviations.

## Discussion

To conclude, we have introduced a method that uses Gaussian processes to infer first and second derivatives from time-series data. In tests, our approach is more accurate than others (Fig. 1), but has several advantages: it systematically estimates errors, both for the regression and the inferred derivatives; it allows interpolation with the corresponding error estimation (Gaussian processes were developed for interpolation [7]); it allows sampling of the latent function underlying the data and so can be used to estimate errors in any statistic of that function by calculating the statistic for the samples; and it can apply inference to any number of experimental replicates.

For fitting growth curves, several alternatives exist [3, 14, 15, 16], which, although mostly focusing on parametric approaches, do allow spline fitting [3] and polynomial regression [14, 16]. Both approaches have been criticised, being sensitive to outliers and potentially having systematic biases [5], and can be out-performed by our algorithm (Fig. 1). Further, our software performs inference using all replicates, can infer second derivatives, and rigorously estimates errors. Where error estimation in summary statistics has been addressed [3], bootstrapping of the data is used. This approach is perhaps less suited for time-series data than ours of sampling latent functions to calculate summary statistics because it leads to some randomly chosen data points being weighted more than others when generating sample fits.

Like any Bayesian method, prior information on bounds for the hyperparameters of the covariance function (the parameters determining the behaviour of the covariance function and optimised by the fitting algorithm) can affect the inference, although these bounds can typically be set so that the best-fit values are far from the bounds. In particular, how closely the latent function follows the data depends both on its flexibility and on the size of the measurement noise. An outlier can be followed if the flexibility is high or if the measurement noise is low. When there is not sufficient data, the algorithm, rightly in our opinion, requires prior information to make this choice. Alternative methods also require prior specification, such as the degree of smoothness in fitting with either splines or a local polynomial method, like LOESS. For a particular type of data, the bounds typically need to be set once allowing high throughput analyses.

Our algorithm is coded in the free computer language Python and can be run both from scripts and a platform-independent GUI.

## Methods

### Overview

A Gaussian process is a collection of random variables for which any subset has a joint Gaussian distribution [7]. To use a Gaussian process to fit a time-series, a random variable is associated with each time point for which there is a measurement. These random variables are characterised by their covariance (the mean is without loss of generality set to zero). The choice of covariance determines a family of functions, and the latent function that underlies the data is assumed to come from this family.

A sample from a Gaussian process (a sample from each random variable associated with a time point) will generate a sample of a function from the family specified by the covariance matrix. For example, if each random variable does not covary with any other (the covariance matrix is the identity matrix), then the functions generated by sampling from the Gaussian process will randomly jump back and forth around zero. If each random variable covaries equally with every other random variable (each entry of the covariance matrix is equal and positive), then the functions sampled will be straight horizontal lines starting at the value sampled for the random variable associated with the first time point. More interestingly, if the covariance for any particular random variable is positive for those random variables whose time points are close in time and tending to zero for random variables far away in time, then the functions generated vary but do so smoothly.

To use Gaussian processes in regression, we must first make a choice of a covariance function. We typically use a squared exponential covariance function, which is a common choice because it is relatively unrestrictive 7, but we have also implemented a neural network covariance function. This covariance function can generate sigmoidal-like curves, which are common in biology, but usually was more sensitive to prior information on the bounds for the hyperparameters.

Given a covariance function and a data set, we must appropriately choose the parameters that specify the covariance function (referred to as the hyperparameters). Typically this choice is made by maximising the marginal likelihood, where the marginalisation is made by integrating over all possible latent functions [7]. Once the parameters of the covariance function have been determined, we can sample latent functions given the data because the probability distribution for the latent functions is Gaussian. Each sampled latent function is a possible function that could underlie the data, and the mean latent function gives the best-fit. The standard deviation of the latent function distribution gives an estimation of the error in the inference at each time point.

Further, we can both infer the time-derivatives. The time-derivative of a Gaussian process is also a Gaussian process [7], and so by fitting the data with a Gaussian process we can also estimate time-derivatives. Errors in fitting are automatically carried through to the errors in inferring time-derivatives.

### Using a Gaussian process to fit time-series data

In the following, we will denote a Gaussian distribution with mean *μ* and covariance matrix Σ as 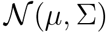 and use the notation of Rasmussen and Williams [7] as much as possible.

For *n* data points *y_i_* at inputs *x_i_* (each *x_i_* is a time for a growth curve), we denote the underlying latent function as *f(x)*. We define a covariance matrix *k(x,x′)*, which has an explicit (although here not written) dependence on hyperparameters θ, and obeys

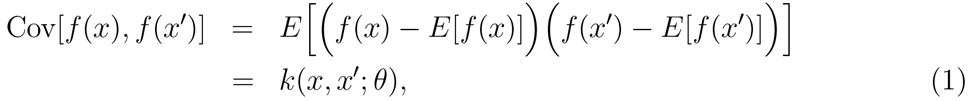

where the expectations are taken over distribution of latent functions (samples of *f(x)*) and θ denotes the covariance function,s hyperparameters.

With Eq. 1, the prior probability distribution is a Gaussian process over the functions *f(X)*, where were we write *X* for the inputs *x_i_* such that

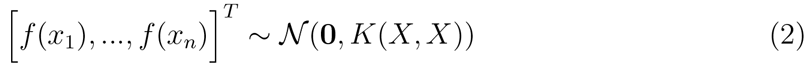

where *K(X,X)* is the n × n matrix with components *k(x_i_,x_j_)*. Given the dependence of *k(x,x′θ)* on the hyperparameters *θ*, we have

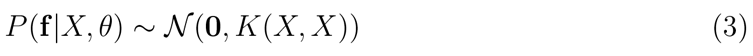

as the prior distribution and writing f for [*f(x_1_),…, f(x_n_)*].

To infer the hyperparameters given the data, we consider the likelihood *P(y|9,X)*, which, more correctly, is a marginal likelihood

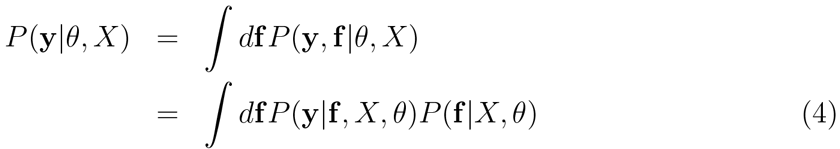

where the marginalisation is over all choices of the latent function f evaluated at *X*.

If we assume that for all *y_i_, y_i_ = f(x_i_) + ∈_i_* where each *∈_i_* is an independent Gaussian variable with zero mean and a standard deviation of σ, then

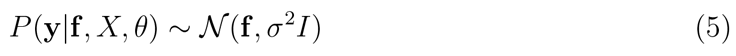

where *I* is the n × n identity matrix. Eqs. 3 and 5 imply that the marginal likelihood is also Gaussian:

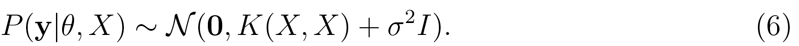

We use a maximum-likelihood method to find the hyperparameters and maximize the marginal likelihood (Eq. 6). We have two hyperparameters for the squared exponential covariance function and the parameter, σ, which characterises the measurement error. We assume a bounded, uniform prior probability for each of these hyperparameters and use the Broyden-Fletcher-Goldfarb-Shanno algorithm [4] to find their optimum values. Although one optimisation run from random initial choices of the hyperparameters is usually sufficient, choosing the best from multiple runs can be better to reduce the algorithm finding local maxima.

### Making predictions

Given the optimum choice of the hyperparameters, we would like to generate sample latent functions at points *X^*^*, which to include the possibility of interpolation need not be the same as X, by sampling from *P*(**f**^*^|*X*, y,θ,*X^*^*). Using Eq. 6 and that the distribution of the latent function evaluated at *X^*^* is also Gaussian, we can write the joint probability of **y** and **f**^*^ as [7]:

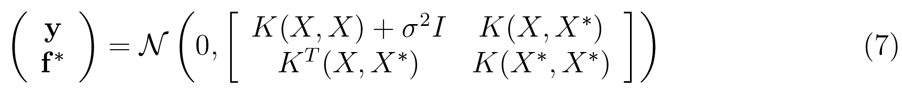

where *K(X,X^*^)* is the n × n^*^ matrix with components 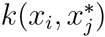.

Conditioning Eq. 7 on the data **y**, standard results for Gaussian distributions [7] give that the probability distribution *P*(**f**^*^|*X*, y,θ,*X^*^*) is also Gaussian with mean 
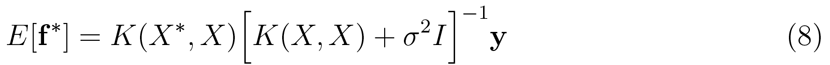

and covariance matrix 
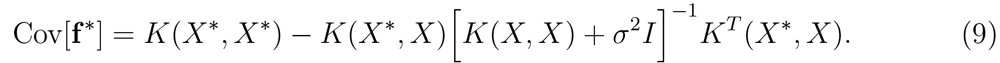

We use Eqs. 8 and 9 to sample f^*^.

### Inferring the first and second time-derivatives

To determine the time-derivative of the data, we use that the derivative of a Gaussian process is another Gaussian process [7]. We can therefore adapt standard techniques for Gaussian process to allow time-derivatives to be sampled too.

Building on the work of Boyle [17], we let *g(x)* and *h(x)* be the first and second derivatives with respect to *x* of the latent function *f(x)*. If *f(x)* is a Gaussian process then so are both *g(x)* and *h(x)*. Writing *∂d*_1_ and *∂*_2_ for the partial derivatives with respect to the first and second arguments of a bivariate function, we have

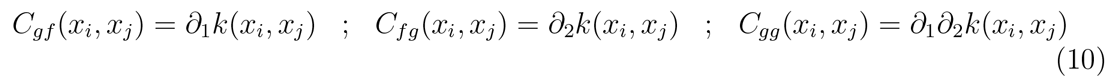

and that

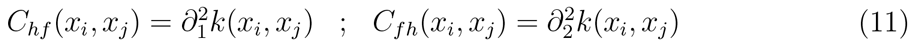

as well as

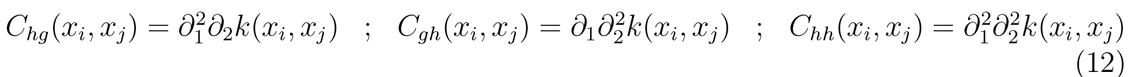

following [18].

Consequently the joint probability distribution for **y** and **f**^*^, **g**^*^, and **h**^*^ evaluated at points *X*^*^ is again Gaussian (cf. Eq. 7):

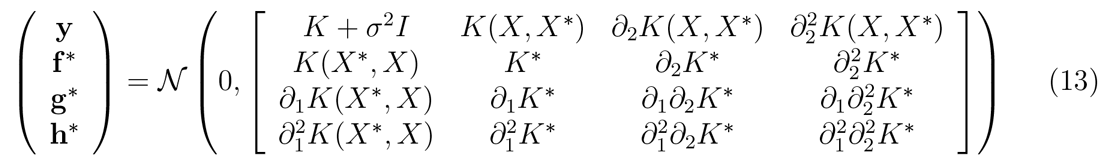

where we write *K = K(X,X)* and *K^*^ = K(X^*^,X^*^)* for clarity.

The covariance function is by definition symmetric: *k(x_i_,x_j_)* = *k(x_j_,x_i_)* (Eq. 1). Therefore 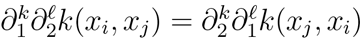 and so

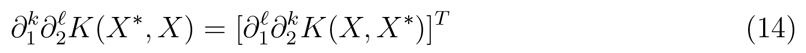

for all positive integers *k* and *l*. Consequently, the covariance matrix in Eq. 13 is also symmetric.

Conditioning on y now gives that the distribution *P*(**f**^*^, **g**^*^, **h**^*^|*X*, y,θ,*X^*^*) is Gaussian with mean

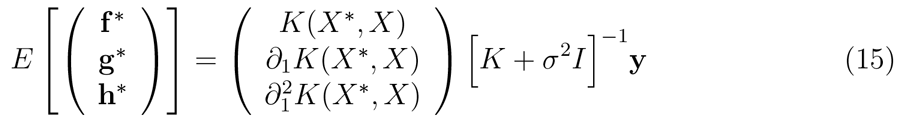

and covariance matrix

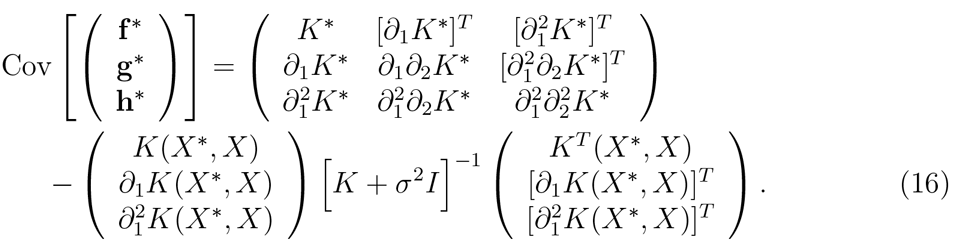

Eq. 16 includes Eq. 9 and shows that

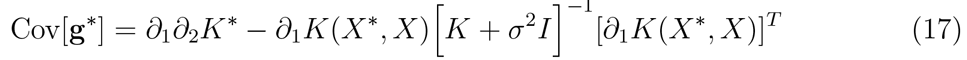

which gives the error in the estimate of the first derivative [17]. Similarly,

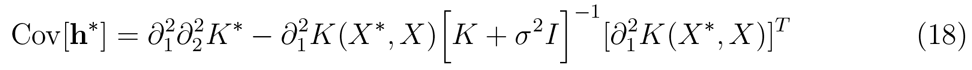
is the error in estimating the second derivative.

### Using an empirically estimated measurement error

Typically we should use a Gaussian process when we expect that the measurement errors in the data are independent and identically distributed with a Gaussian distribution of mean zero. When this assumption does not appear to be true, we have an alternative implementation where we empirically estimate the measurement error but to do so we require multiple experimental replicates. We calculate the variance across all replicates at each time point and then smooth over time (with a Gaussian filter with a width of 10% of the total time of the experiment, but the exact choice in not important).

To make predictions, we replace the identity matrix, *I*, in Eqs. 6, 15, and 16 by a diagonal matrix with the empirical measurement errors on the diagonal.

### Estimating the growth characteristics

From the growth curve, we estimate the maximum growth rate as the maximum time-derivative of the logarithm of the growth curve [2]:

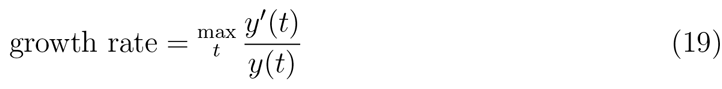

where we denote the growth curve as *y(t)*. The doubling time is ln(2) times the inverse of the growth rate. We define the lag time as the intercept of the line parallel to the time axis that passes through the initial OD, *y*(0), and the tangent to the logarithm of the growth curve from the point on the growth curve with maximum growth rate (a standard choice [2]). If this point of maximum growth rate is at *t* = *t*^*^, then

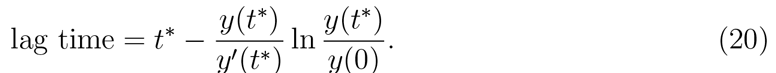

For each characteristic, we can estimate measurement errors through sampling latent growth curves consistent with the data.

### Implementation and GUI

The code for our algorithm is freely available and written in Python 3 using NumPy [19], SciPy (Jones, Oliphant, Peterson, *et al.*), Matplotlib [20], and the Pandas data analysis library (all available via the free Anaconda package) and is compatible with Microsoft,s Excel. We give an example script and data set and have written a GUI that runs on Windows, OS X, and Linux.

The software and instructions for its use are at http://swainlab.bio.ed.ac.uk/software/fitderiv

### Experimental methods

Data for Fig. 2A was gathered using Tecan Infinity M200 plate reader and a BY4741 strain of *S. cerevisiae* growing in synthetic complete media supplemented with 0.4% glucose and 1% galactose at 30°C, following an established protocol [21]. OD was measured at an absorbance wavelength of 595 nm every 11.4 minutes.

Data for Fig. 2B was gathered using a Spectrostar Omega microplate reader and a BW25113 strain of *E. coli* growing in MM9 (sodium-sodium instead of sodium-potassium) media with 0.1% glucose and 1106 mOsm sucrose at 37°C. OD was measured at an absorbance wavelength of 600 nm every 7.5 minutes.

**Table 1.** Ranges of hyperparameters used for the examples. For the squared exponential covariance function, the hyperparameters determine the amplitude of the variation in the latent function, its flexibility, and the magnitude of the measurement error; for the neural network covariance function, the hyperparameters determine the initial *y*-value of the latent function, its flexibility, and the magnitude of the measurement error.

| Figure | Covariance function | Hyperparameter | Lower bound | Upper bound |
| --- | --- | --- | --- | --- |
| 1A | squared exponential | 0 | $10^{-5}$ | $10^5$ |
| | | 1 | $10^{-3}$ | $10^2$ |
| | | 2 | $10^{-5}$ | $10^2$ |
| | neural network | 0 | $10^{-1}$ | $10^5$ |
| | | 1 | $10^3$ | $10^3$ |
| | | 2 | $10^{-6}$ | $10^2$ |
| 1B | squared exponential | 0 | $10^{-5}$ | $10^5$ |
| | | 1 | $10^{-3}$ | $10^4$ |
| | | 2 | $10^{-5}$ | $10^2$ |
| | neural network | 0 | $10^{-1}$ | $10^5$ |
| | | 1 | $10^1$ | $10^{2.5}$ |
| | | 2 | $10^{-6}$ | $10^2$ |
| 2A & 2B | squared exponential | 0 | $10^{-5}$ | $10^5$ |
| | | 1 | $10^{-6}$ | $10^2$ |
| | | 2 | $10^{-5}$ | $10^2$ |
| 3A | squared exponential | 0 | $10^{-5}$ | $10^5$ |
| | | 1 | $10^{-6}$ | $10^2$ |
| | | 2 | $10^{-5}$ | $10^0$ |
| 3B | neural network | 0 | $10^{-1}$ | $10^5$ |
| | | 1 | $10^{-4}$ | $10^{-1}$ |
| | | 2 | $10^{-6}$ | $10^2$ |

Data for Fig. 3A is from [11].

Data for Fig. 3B was gathered using a custom spinning disk confocal microscope for 20mins in 20s time steps with 50ms exposure time per focal plane. Spindle pole bodies were labelled with Spc42-Cerulean. An image stack of 30 *z*-planes with 300nm step size was gathered for each time point to allow the position of the spindle poles to be fitted to 3-D Gaussian distributions and tracked in time. Imaging, fitting and tracking followed an established protocol [12].

Data generated in this work is available at http://dx.doi.org/10.7488/ds/1405.

## Acknowledgements

We thank Guido Sanguinetti and Nacho Molina for advice on Gaussian processes, Ryan Morris and Cait MacPhee for providing the data in Fig. 3A, and our funders: a BBSRC iCASE award (KS), the Wellcome Trust and Conacyt (LFMG), and the CIHR (AL & JV). PSS, IBNC, and TP gratefully acknowledge the support of the UK Research Councils Synthetic Biology for Growth programme and are members of a BBSRC/EPSRC/MRC funded Synthetic Biology Research Centre. The authors have no competing financial
interests.

## Author contributions

PSS developed the algorithm and, with KS & AL, the code; KS, AL, LFM, IBNC, & JV generated the data; TP & PSS wrote the manuscript.

